# A Computational Model for the Evaluation of Complement System Regulation under Homeostasis, Disease, and Drug Intervention

**DOI:** 10.1101/225029

**Authors:** Nehemiah Zewde, Dimitrios Morikis

## Abstract

**Highlights:**

- Computational model describing dynamics of complement system activation pathways
- Complement dysregulation leads to deviation from homeostasis and to inflammatory diseases
- Model identifies biomarkers to quantify the effects of complement dysregulation
- Known drugs restore impaired dynamics of complement biomarkers under dysregulation
- Disease-specific models are suitable for diagnosis and patient-specific drug treatment

*Abstract:* The complement system is a part of innate immunity that rapidly removes invading pathogens and impaired host-cells. Activation of the complement system is balanced under homeostasis by regulators that protect healthy host-cells. Impairment of complement regulators tilts the balance, favoring activation and propagation that leads to inflammatory diseases. The most potent regulator of the complement system is Factor H (FH), and its impairment induces improper complement activation that leads to inflammatory diseases, such as atypical hemolytic uremic syndrome and age related macular degeneration. To understand the dynamics involved in the pivotal balance between activation and regulation, we have developed a comprehensive computational model of the alternative and classical pathways of the complement system. The model is composed of 290 ordinary differential equations with 142 kinetic parameters that describe the state of complement system under homeostasis and disorder through FH impairment. We have evaluated the state of the system by generating concentration-time profiles for the biomarkers C3, C3a-desArg, C5, C5a-desArg, Factor B (FB), Ba, Bb, and fC5b-9 that are influenced by complement dysregulation. We show that FH-mediated disorder induces substantial levels of complement activation compared to homeostasis, by generating reduced levels of C3 and FB, and to a lesser extent C5, and elevated levels of C3a-desArg, Ba, Bb, C5a-desArg, and fC5b-9. These trends are consistent with clinically observed biomarkers associated with complement-mediated diseases. Furthermore, we introduced therapy states by modeling known drugs of the complement system, a compstatin variant (C3 inhibitor) and eculizumab (a C5 inhibitor). Compstatin demonstrates strong restorative effects for early-stage biomarkers, such as C3a-desArg, FB, Ba, and Bb, and milder restorative effects for late-stage biomarkers, such as C5a-desArg and fC5b-9, whereas eculizumab has strong restorative effects on late-stage biomarkers, and negligible effects on early-stage biomarkers. These results highlight the need for patient-specific therapies that target early complement activation at the C3 level, or late-stage propagation of the terminal cascade at the C5 level, depending on the specific FH-mediated disease and the manifestations of a patient’s genetic profile in complement regulatory function.

---

**Nomenclature**

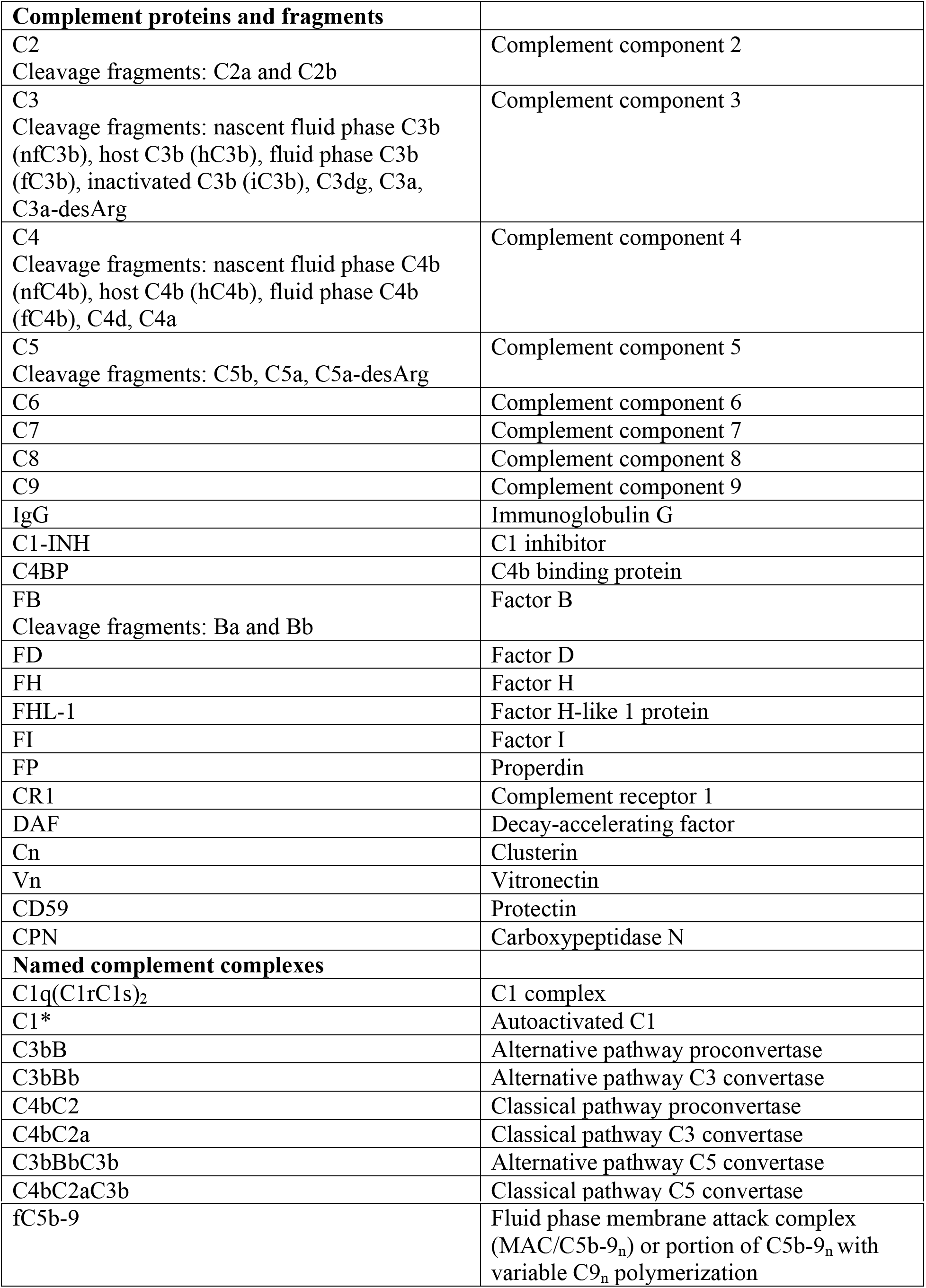

|  |  |
| --- | --- |
| **Complement proteins and fragments** |  |
| C2<br>Cleavage fragments: C2a and C2b | Complement component 2 |
| C3<br>Cleavage fragments: nascent fluid phase C3b (nfC3b), host C3b (hC3b), fluid phase C3b (fC3b), inactivated C3b (iC3b), C3dg, C3a, C3a-desArg | Complement component 3 |
| C4<br>Cleavage fragments: nascent fluid phase C4b (nfC4b), host C4b (hC4b), fluid phase C4b (fC4b), C4d, C4a | Complement component 4 |
| C5<br>Cleavage fragments: C5b, C5a, C5a-desArg | Complement component 5 |
| C6 | Complement component 6 |
| C7 | Complement component 7 |
| C8 | Complement component 8 |
| C9 | Complement component 9 |
| IgG | Immunoglobulin G |
| C1-INH | C1 inhibitor |
| C4BP | C4b binding protein |
| FB<br>Cleavage fragments: Ba and Bb | Factor B |
| FD | Factor D |
| FH | Factor H |
| FHL-1 | Factor H-like 1 protein |
| FI | Factor I |
| FP | Properdin |
| CR1 | Complement receptor 1 |
| DAF | Decay-accelerating factor |
| Cn | Clusterin |
| Vn | Vitronectin |
| CD59 | Protectin |
| CPN | Carboxypeptidase N |
| **Named complement complexes** |  |
| C1q(C1rC1s)<sub>2</sub> | C1 complex |
| C1* | Autoactivated C1 |
| C3bB | Alternative pathway proconvertase |
| C3bBb | Alternative pathway C3 convertase |
| C4bC2 | Classical pathway proconvertase |
| C4bC2a | Classical pathway C3 convertase |
| C3bBbC3b | Alternative pathway C5 convertase |
| C4bC2aC3b | Classical pathway C5 convertase |
| fC5b-9 | Fluid phase membrane attack complex (MAC/C5b-9<sub>n</sub>) or portion of C5b-9<sub>n</sub> with variable C9<sub>n</sub> polymerization |

## 1. Introduction

A pivotal arm of innate immunity, the complement system is comprised of proteins present in both plasma and cell membranes that work in concert to mediate immune responses against invading pathogens and altered host cells, while at the same time shielding healthy host cells from destruction [1,2]. The coordination of the complement system is induced through three pathways known as alternative, classical, and lectin. The alternative pathway is constitutively activated in plasma through the so-called tick-over mechanism of complement protein C3, while the latter two pathways, classical and lectin, are initiated on microbes or apoptotic/necrotic cells by their respective pattern recognition molecules [3]. Similar to alternative pathway, the classical pathway can also be induced in the fluid phase through the spontaneous intramolecular activation of complement complex C1 [4]. Central points of convergence for all three pathways of complement activation and propagation are complement proteins C3 and C5, whose cleavage fragments C3a/C3b and C5a/C5b, respectively, results in inflammatory responses, opsonophagocytosis, and formation of the membrane attack complex (MAC/C5b-9_n_) [5,6]. The fragments C3a and C5a are anaphylatoxins that mediate activation of immune cells, such as macrophages and T cells, while C3b is an opsonin that tags foreign or altered cells for recognition and elimination by phagocytes [5,6]. Lastly, the MAC complex penetrates lipid bilayers and induces lysis through osmotic effects [7]. Propagation of complement activation occurs through cleavage of C3 or C5 by convertases, which are complexes of cleavage products of C3, C4, and proteases Factor B (FB) and C2 [8–11].

The complexity of the complement system comes from the number of proteins and their networks of interactions involved in all three pathways. However, recent mathematical models targeting complement pathways have been developed to aid our understanding in the dynamics involved in mediating immune response through activation and regulation of complement species [12–15]. To further these efforts, we previously developed a comprehensive model of the alternative pathway divided into four modules: (i) initiation (fluid phase); (ii) amplification (surfaces); (iii) termination (pathogen); and (iv) regulation (host cell and fluid phase) [16]. Based on these four modules we generated a system of 107 ordinary differential equations (ODEs) and 74 kinetic parameters. Here we present an updated and refined alternative pathway model of the complement system that takes into account dimerization states of complement fragments C3b and C4b, additional C3 and C5 convertases, IgG interactions with the C3b, and regulation of anaphylatoxins C3a and C5a. Relative to our first model [16], this new model also contains fluid phase activation and propagation of the classical pathway, owed to the spontaneous autoactivation of C1. With these additions, our updated and refined model is now composed of 290 ordinary differential equations and 142 kinetic parameters.

The cascade of reactions “innate” to the complement system is equally supplemented with interactions that protect healthy host cells from destruction, under homeostasis. Recent studies highlight a number of diseases that are associated with under-regulation/over-activation of the alternative pathway due to mutations that target regulators (e.g. Factor H, FH) or propagators (convertases) of the alternative pathway [17–22]. Diseases such as such as atypical hemolytic uremic syndrome (aHUS), age related macular degeneration (AMD), C3 glomerulonephritis (C3GN), and dense-deposit disease (DDD) are associated with overly active alternative pathway [18–21, 23–27]. To better understand the dynamics of these disorders at the complement level, we have developed a generic disease state by targeting the most potent complement regulator FH. This regulator is chosen because alternative pathway-mediated disorders, mentioned above, implicate FH dysfunctionality, owed to altered effective concentrations or mutations that impair regulatory capabilities. In addition to modeling the disease state, we have also added to two therapy states by incorporating complement inhibitors, compstatin, targeting C3 [28–32], and eculizumab, targeting C5 [33–35], and compared their relative effects on regulating an overactivated complement system under FH regulatory impairment. Furthermore, we implemented different doses of compstatin and eculizumab to investigate their relative roles on mediating early and late stage complement biomarkers. In summary, we have generated four states of alternative and classical pathways with time profiles describing concentration levels of biomarkers C3, C3a-desArg, C5, C5a-desArg, FB and its fragments Ba and Bb, and fC5b-9 (a general term for fluid phase MAC/C5b-9) that are associated with alternative pathway dysregulation.

## 2. Methods

### 2.1 Mathematical model

Our mathematical model is based on the biochemical reactions shown in Fig 1. The biochemical reactions are generated from critical reading of papers with experimental and clinical data, published in the scientific literature. Using the cascade of biochemical reactions in the classical and alternative pathway, we generated a system of 290 ODEs with 142 kinetic parameters. Similar to our previous model [16], we organized the biochemical reactions and equations into modules that describe complement system: (i) initiation, Eqs S1–S55 of the Supplementary Material; (ii) amplification, Eqs S56–S77 of the Supplementary Material; (iii) termination, Eqs S78–S110 of the Supplementary Material; (iv) regulation Eqs S111–S256 of the Supplementary Material; and (v) complement proteins in fluid phase and derived from bound components on host cells, Eqs S257-S290 of the Supplementary Material. In addition, we added two more modules that describe complement therapeutic states brought by the actions of known complement inhibitors compstatin, Eqs S291–S293 of the Supplementary Material, and eculizumab, Eqs S294–S296 of the Supplementary Material. As mentioned above, compstatin targets complement component C3 and eculizumab targets complement component C5. In this study, we use kinetic parameters for the compstatin analog with sequence Ac-I[CVWQDWGAHRC]T-NH_2_ (brackets denote disulfide bridge cyclization) from Ref [32].

**Fig 1.**
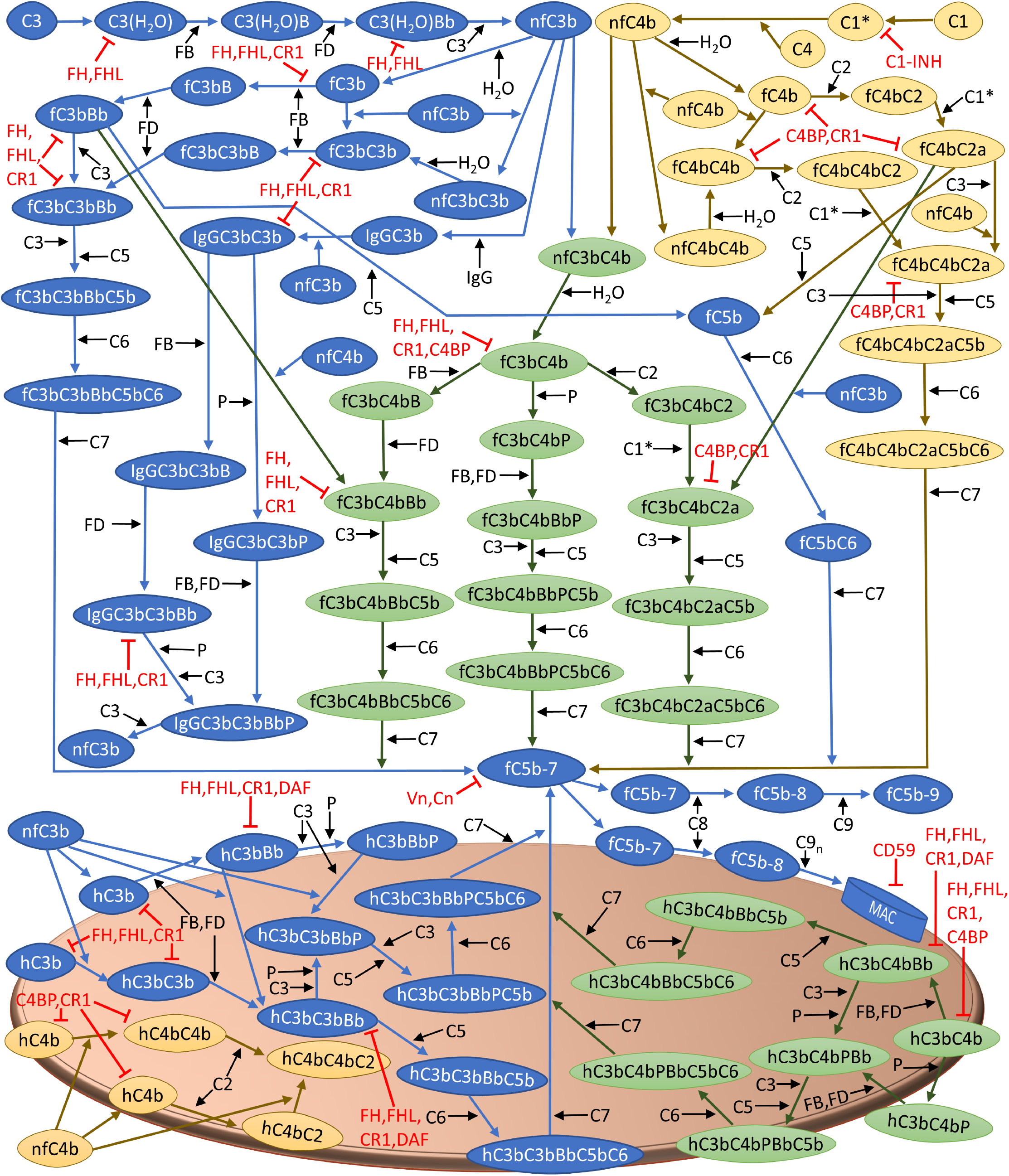
Biochemical reactions originating from fluid phase activation of the alternative and classical pathways. A complete explanation of the various steps, grouped in modules of initiation (fluid phase), amplification, termination, and regulation can be found in Section 2.2 Biochemical Model of Methods. Initiation of the alternative pathway (blue) is induced through the tick-over reaction of C3, while classical pathway (yellow) is activated through the spontaneous intramolecular activation of complement complex C1. The tick-over of C3 in the alternative pathway forms the C3 convertase, C3(H_2_O)Bb, that cleaves C3 to generate nascent fluid phase C3b (nfC3b) and C3a, whereas activation of C1 cleaves C4 into nascent fluid phase C4b (nfC4b) and C4a. Subsequently, nfC3b and nfC4b can bind to host cells, form dimers such as C3bC4b (green), C3bC3b, and C4bC4b, or generate fluid phase C3b and C4b (fC3b and fC4b). Formation of fC3b and fC4b initiates a cascade of reactions that forms C3 and C5 convertases (fC3bBb and fC4bC2a). These enzymes (fC3bBb and fC4bC2a) further cleave plasma C3 to produce more nfC3b and C3a. Furthermore, fC3bBb and fC4bC2a can also cleave C5 to generate C5a and C5b. Continued propagation of complement through C5b leads to C5b-7 that can also attach to host cells and initiate a cascade of reactions to form MAC (C5b-9_n_). However, healthy host cells are shielded from complement attack due to fluid phase and surface bound complement regulators shown in red. Regulators FH, FHL-1, C4BP, C1-INH, CR1, and DAF provide early phase checkpoints for complement activation and propagation, while late-step regulation of the terminal cascade is provided by fluid regulators Vn/Cn and surface bound regulator CD59.

Perturbation that leads to regulatory disorder is modeled by reducing FH concentration and kinetics by an order of magnitude as shown in Tables S1 and S2 of Supplementary Material, respectively. The biochemical reactions are converted to a system of ODEs, describing the mass balance for each complement protein, protein fragment, or protein complex. Enzymatic reactions are based on Michaelis–Menten kinetics and substrate competitions for the same enzyme are considered for complement species such as C3 and C5 convertases. Equations are solved using the ode15s solver of Matlab (Mathworks, Natick, MA).

Methods for parameter estimations or calculations, when experimental values are not available (Tables S1 and S2 of Supplementary Material), and multi-parametric sensitivity analysis (MPSA) for global sensitivity are described in our previous paper [16]. Briefly, for MPSA we first selected kinetic parameters involved in our biochemical reactions of alternative and classical pathways. Then we generated ranges for each parameter to be large enough to cover all feasible variations. For each parameter set, we calculated the sum of squared errors between the observed and perturbed system output values. We used the results of this calculation to determine if the chosen set of parameters is acceptable or unacceptable. The cumulative frequency was calculated for both acceptable and unacceptable cases and was used to evaluate the sensitivity of each parameter using Kolmogorov-Smirnov statistic. Parameter ranges used in multi-parametric sensitivity analysis can be found in Table S3 of Supplementary Material.

For unknown parameters, we assumed the kinetic rate constants to be the same as in those of structurally or functionally homologous proteins with known experimental parameter values. For instance, the kinetic parameters for C3(H_2_O) and FB interaction are not known, whereas those for C3b and FB interaction are known. Since C3(H_2_O) is known to be a C3b-like molecule that functions similarly to C3b by forming alternative pathway proconvertases and C3 convertases, we assumed C3(H_2_O) to have the same kinetic parameters as C3b. In addition, since the convertases C3(H_2_O)Bb and C3bBb decay with similar rates, 9.0×10^−3^ s^-1^ and 7.7×10^−3^ s^-1^, respectively [36], we assumed the rate of decay induced by FH on C3bBb and on C3(H_2_O)Bb to be the same. This assumption is consistent with similar biochemical interaction between FH and serine protease, Bb, that is present on both convertases (C3(H_2_O)Bb, C3bBb). Lastly, due to the lack of kinetic data on the decay rates induced by complement regulators such as decay-accelerating factor (DAF), complement receptor 1 (CR1), and C4b-binding protein (C4BP), we assumed they will also have the same decay accelerating rate as FH because these regulators function in a FH-like manner by either inhibiting or deactivating C3/C5 convertases.

### 2.2 Biochemical Model

Below, we will describe the complement system pathways of Fig 1. These pathways entail activation, propagation, and regulation of the complement system. The pathways describe interactions of complement proteins, fragments, and complexes that operate entirely in the fluid phase, as well as those that bind to cell surfaces, undergoing further processing, and returning products to fluid phase. The characteristics of the surface of a red blood cell are used in the model [16].

#### 2.2.1 Initiation (fluid phase)

The alternative pathway is spontaneously activated in plasma through a process known as the tick-over reaction, and involves the spontaneous hydrolysis of an internal thioester bond in C3. This is an irreversible step that produces a C3b-like molecule, C3(H_2_O), with its internal thioester bond hydrolyzed. The complement protease FB associates with C3(H_2_O) to form a proconvertase, C3(H_2_O)B. Subsequently, enzymatic cleavage of C3(H_2_O)-bound FB by Factor D (FD) follows to form the initial convertase of the alternative pathway, C3(H_2_O)Bb. This shortlived enzyme can cleave C3 to form C3a and nascent fluid phase C3b (nfC3b). While C3a is anaphaytoxin [5], nfC3b has an exposed highly-reactive internal thioester bond that is capable of indiscriminately binding to different surfaces via covalent attachment, or forming fluid phase C3b (fC3b) by interacting with water [37–39]. The fC3b fragment cannot covalently attach to cells but can still propagate the alternative pathway by associating with FB and following the same cascade of reactions as C3(H_2_O), forming form the fluid C3/C5 convertase, C3bBb [36,40].

Similar to the alternative pathway, the classical pathway also initiates in the fluid phase through the spontaneous autoactivation of component complex C1 [4]. This leads to the formation of activated C1 (denoted C1*) that cleaves C4. This process leads to the formation of C4a and nascent fluid phase C4b (nfC4b). Similar to nfC3b, nascent fluid C4b can covalently attach to nearby surface or form fluid phase C4b (fC4b) by interacting with water [41]. Complement C2 then interacts with C4b to form the proconvertase of the classical pathway, C4bC2. Similar to C4, this complex is also activated by C1* to form the C3/C5 convertase, C4bC2a.

In addition to the covalent bonds formed between nfC3b/nfC4b to cell membranes, covalent linkage can occur between C3b and C4b fragments, forming C3bC4b, C3bC3b, or C4bC4b complexes. These complexes can associate with FB or C2 to form C3/C5 convertases, fC4bC3bBb, fC3bC3bBb, fC3bC4bC2a, and fC4bC4bC2a [9–11]. Furthermore, C3b dimers, C3bC3b, can also covalently attach with immunoglobulin G, IgG, to form IgGC3bC3b [8]. This complex functions as a C3 convertase that cleaves C3 into C3a and nfC3b.

#### 2.2.2. Amplification

Once nfC3b attaches covalently to a host surface, a cascade of reactions ensues where FB is recruited first, followed by FD cleavage to yield the surface bound C3/C5 convertase, C3bBb. Since this enzyme is rather unstable, the only positive regulator of the alternative pathway, known as properdin, binds and extend the half-life of C3bBb by 10-fold [42,43]. Similar to the propagation step in the fluid phase, formation of dimers associated with C3b or C4b occurs on the surface of host cells. These dimers, C3bC3b/C3bC4b, can associate with either FB, followed by activation step with FD to form the C3/C5 convertases (C3bC3bBb/C4bC3bBb). However, unlike C3bC3b/C3bC4b dimers in the fluid phase that can also bind to C2, the surface activation mechanism for C1* is not added since it requires antigens to anchor to the surface first [44–46]. Here, host cells are under normal conditions while the impairment in our model only applies to FH in the fluid phase. Lastly, C3b dimers can also associate with properdin to form stabilized C3/C5 convertase that enhances complement activation within the vicinity of this convertase.

#### 2.2.3. Termination

Cleavage of complement C5 by C3/C5 convertases such as fC3bBb or fC4bC2a, leads to the formation of C5a and C5b. Like C3a, C5a is a mediator of inflammation [5]. C5b binds to C6 and subsequently to C7 and forms C5b-7. This complex can insert itself into the lipid bilayer but also can form inactive micelles in the absence of cell membranes [47,48]. Furthermore, C5b-7 binds to C8, followed by C9 to form fluid phase MACs (fC5b-9). However, if C5b-7 attaches to a cell, C8 binding occurs, followed by the attachments of multiple C9 molecules to from a MAC. Presence of MACs compromise the integrity of cell membranes and induce cell death. In addition to fC3bBb/fC4bC2a cleaving C5, dimerized forms such as fC4bC3bBb or fC3bC4bC2a can also cleave C5 but C5b remains loosely bound to the convertase while C5a is released in the fluid phase. When C5b is still bound to the convertase, it is capable of binding to C6. Complement C7 then binds and subsequently leads to C5b-7 complex dissociating from the C3/C5 convertase.

#### 2.2.4. Regulation

Multiple check-points are in place to ensure proper regulation of the complement system. FH and Factor H-like 1 (FHL-1) proteins are primarily regulators of the alternative pathway that can act in both fluid and surface phases, while C1 inhibitor (C1-INH) and C4BP are the primary regulators for the classical pathway in both phases (fluid and surface). These regulators of complement act on the early stages of the activation by targeting C1* (by C1-INH), or C3b and C4b (by C4BP) to suppress the propagation step. Inactivation of C3b is mediated through its cleavage to inactivated C3b (iC3b) by Factor I (FI) in conjunction with either FH, FHL-1, or CR1 as cofactors, while C4b is deactivated into C4d by FI with C4BP as its cofactor. Further cleavage of iC3b to C3dg is mediated by FI, using CR1 as a cofactor [1,2]. In addition, other membrane bound regulators such DAF can also act in concert with FH, FHL-1, and C4BP to accelerate the decay of C3/C5 convertases [1,2]. Furthermore, membrane cofactor protein (MCP) is another cell-bound regulator that acts as a cofactor to deactivate C3b/C4b in conjunction with FI. Since MCP is not expressed on erythrocytes, it was not included in the model. Finally, late stage complement propagation in the terminal cascade is tightly controlled through fluid phase regulators such as Vitronectin (Vn) or Clusterin (Cn) that act on fC5b-7, while surface bound regulator CD59 inhibits C9 polymerization [7,49].

#### 2.2.5. FH disorder, with compstatin and eculizumab treatments

Disorders of the alternative pathway, such as C3GN, DDD, AMD, and aHUS all implicate FH impairment through either reduced concentration (type 1 mutation) or binding/regulatory functionality (type 2 mutation) [18–27]. To mimic clinical observations, we generated an FH disorder model by reducing its concentration and association rate constant by one order of magnitude (Tables S1 and S2 of Supplementary Material). After modeling FH disorder, we subsequently generated complement therapeutic states by incorporating into the FH disorder state known complement inhibitors, such as a compstatin family peptide and the monoclonal antibody eculizumab. Compstatin is small peptide that targets complement C3 and inhibits the cleavage of C3 by C3/C5 convertases, [28–30]. Similar in functionality to compstatin, eculizumab targets complement C5 and inhibits the cleavage of C5 by C3/C5 convertases [33–35].

#### 2.2.6. Dose dependence for compstatin and eculizumab

In the initial calculations, the concentrations of compstatin and eculizumab were taken to be 20-fold higher than the concentrations of their respective targets, C3 and C5 (S1 Fig of Supplementary Material). These drug concentrations are used to generate Figs 2–5. In addition, we performed calculations using two more drug concentrations to evaluate the dose-dependent effect on complement dynamics under the disease state induced by FH impairment. Concentrations of compstatin and eculizumab were taken to be one-to-one and 5-fold lower than the concentrations of their respective targets. In total, in the dose-dependent study of compstatin we used concentrations: 1.4×10^−4^M (20-fold higher than C3), 7.1×10^−6^M (1-to-1 to C3) and 1.4× 10^−6^M (5-fold lower than C3). In the dose-dependent study of eculizumab we used concentrations: 7.4×10^−6^M (20-fold higher than C5), 3.7×10^−7^M (1-to-1 to C5) and 7.4×10^−8^M (5-fold lower than C5).

**Fig 2.**
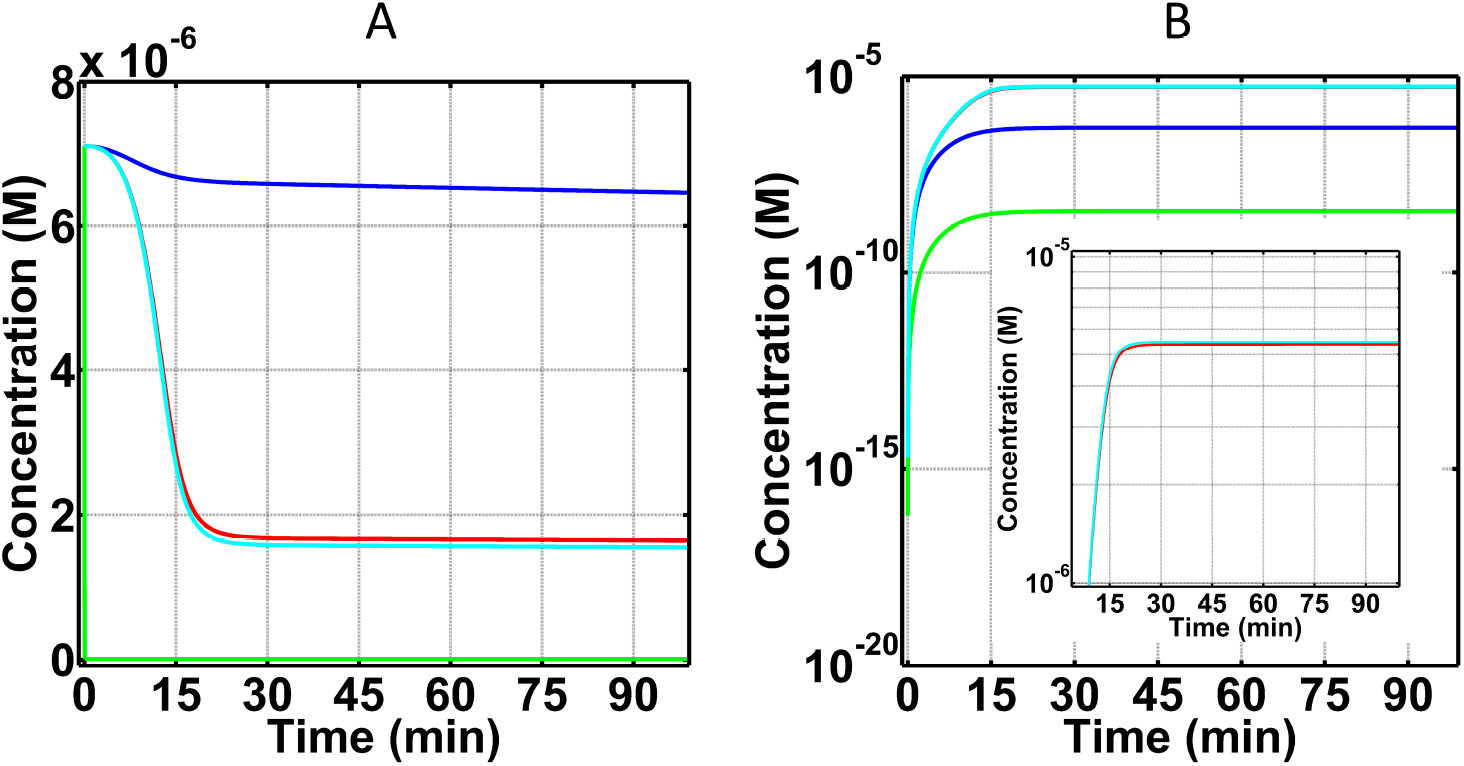
Concentration-time profiles for C3 and C3a-desArg concentrations under four conditions: (i) normal state (blue), (ii) FH disorder state (red), (iii) FH disorder state with compstatin treatment (green), and (iv) FH disorder state with eculizumab treatment (cyan). (A) Consumption of C3 under the four conditions. Plasma C3 is mostly consumed in the FH disorder state, while treatment with eculizumab had small effects on the consumption of C3. Treatment with compstatin leads to the formation of compstatin:C3 complexes, thus leaving small amounts of unbound C3 remain in plasma. (B) Production of C3a-desArg after cleavage of C3. Compstatin shows an over-restoration effect in the FH disorder state. Eculizumab has minor effects on C3 and C3a-desArg in the FH disorder state, with the effect on C3a-desArg being shown with more clarity in the zoomed-in inset of (B).

## 3. Results

### 3.1. Dynamics of complement activation, propagation, regulation, and inhibition

Our study presents the time profiles of the concentrations of complement components for biochemical reactions involved in the alternative and classical pathway of the complement system under four conditions: (i) normal state, corresponding to homeostasis in a healthy person, (ii) FH disorder state, corresponding to alternative pathway dysregulation owed to FH impairment, (iii) FH disorder state with compstatin treatment, and (iv) FH disorder state with eculizumab treatment.

#### 3.1.1. C3 and its cleavage fragment, C3a-desArg, as biomarkers for FH disorder state

We first generated time profiles for the complement system’s key substrate, C3, and its deactivated anaphylatoxin, C3a-desArg, as shown in Figs 2A and 2B. C3 under normal condition has a slow rate of consumption in which C3 levels reach a concentration of 6.6× 10^−6^M within the first 25 minutes from a starting concentration of 7.1 × 10^−6^M (Fig 2A). This change corresponds to a 7% decrease in blood plasma C3 within 25 minutes. The trend of C3 reduction continues but the rate of consumption is significantly reduced after 25 minutes. C3 levels however in FH disorder state and FH disorder state with eculizumab treatment reach a concentration of 1.7× 10^−6^M and 1.6×10^−6^M, respectively, within the first 25 minutes as shown in Fig 2A. (Eculizumab acts on C5 in a downstream step, and therefore has a small effect on C3 concentration.) This change corresponds to a 76% and 77% decrease, respectively, relative to normal condition. In contrast, FH disorder with compstatin treatment left small amounts blood plasma C3 (1.2× 10^−8^M), because C3 forms a new complex with compstatin, CompC3 (Eqs S291-S293 of the Supplementary Material). In other words, this is not owed to C3 being consumed in a biochemical reaction, such as a cleavage process where C3 is converted into nC3b and C3a through the actions of C3/C5 convertases. Rather, through compstatin binding to free C3, a new compstatin:C3 complex is formed. However, for the remaining C3 concentration, not in complex with compstatin, cleavage will follow to form nC3b and C3a. The latter product (C3a) is then rapidly deactivated by carboxypeptidase to form C3a-desArg. Because of this rapid deactivation, C3a-desArg is a better biomarker than C3a. Levels of C3a-desArg are indicators of the degree of complement activation and propagation, and in FH disorder state and FH disorder state with eculizumab treatment, C3a-desArg shows the highest production levels reaching concentrations of 5.4×10^−6^M and 5.5×10^−6^M within 25 minutes, respectively (Fig 2B). In contrast, C3a-desArg levels are about an order of magnitude lower (4.6× 10^−7^M) in the normal state, and about three orders of magnitude lower (3.5×10^−9^M) FH disorder state with compstatin treatment, relative to the FH disorder state and FH disorder state with eculizumab treatment.

#### 3.1.2. C5 and its cleavage fragment, C5a-desArg, as biomarkers for FH disorder state

Terminal component C5, shown in Fig 3A, exhibits minor changes in blood plasma concentrations with the exception being under treatment with eculizumab. All three conditions of normal state, FH disorder state, and FH disorder state with compstatin treatment generate time profiles of C5 where the percent change is < 1% relative to the starting blood plasma concentration of C5 (3.7×10^−7^M). Conversely, treatment with eculizumab significantly reduces blood plasma C5 (1.0×10^−12^M) as shown in Fig 3A. This reduction of C5 is not caused by cleavage, but it is because of the formation of a new species, the eculizumab:C5 complex, EcuC5 in Eqs S294-S296 of the Supplementary Material. However, for the remaining C5 concentration that is not in complex with eculizumab, cleavage will follow to produce C5b and C5a in which C5a is rapidly deactivated by carboxypeptidase to form C5a-desArg. As is the case of C3a/C3a-desArg, this rapid cleavage makes C5a-desArg a better biomarker than C5a. Figure 3B presents the production levels of C5a-desArg in the four states (normal, FH disorder, FH disorder with compstatin treatment, and FH disorder with eculizumab treatment). The highest C5a-desArg levels are produced in FH disorder state, where the concentration of C5a-desArg reaches 5.4×10^−10^M within the first 25 minutes. This upward trajectory continues but the rate of production is significantly reduced after 25 minutes. In contrast, FH disorder state with compstatin treatment produced lower levels of C5a-desArg with concentration 1.2×10^−10^M. This concentration is higher than in the normal state, which produces 2.7× 10^−11^M of C5a-desArg within the same 25 minutes. Lastly, treatment with eculizumab produced the largest difference in C5a-desArg levels, where the concentration reached 3.5×10^−15^M, about five orders of magnitude lower compared to that of FH disorder state C5a-desArg levels (5.4×10^−10^M).

**Fig 3.**
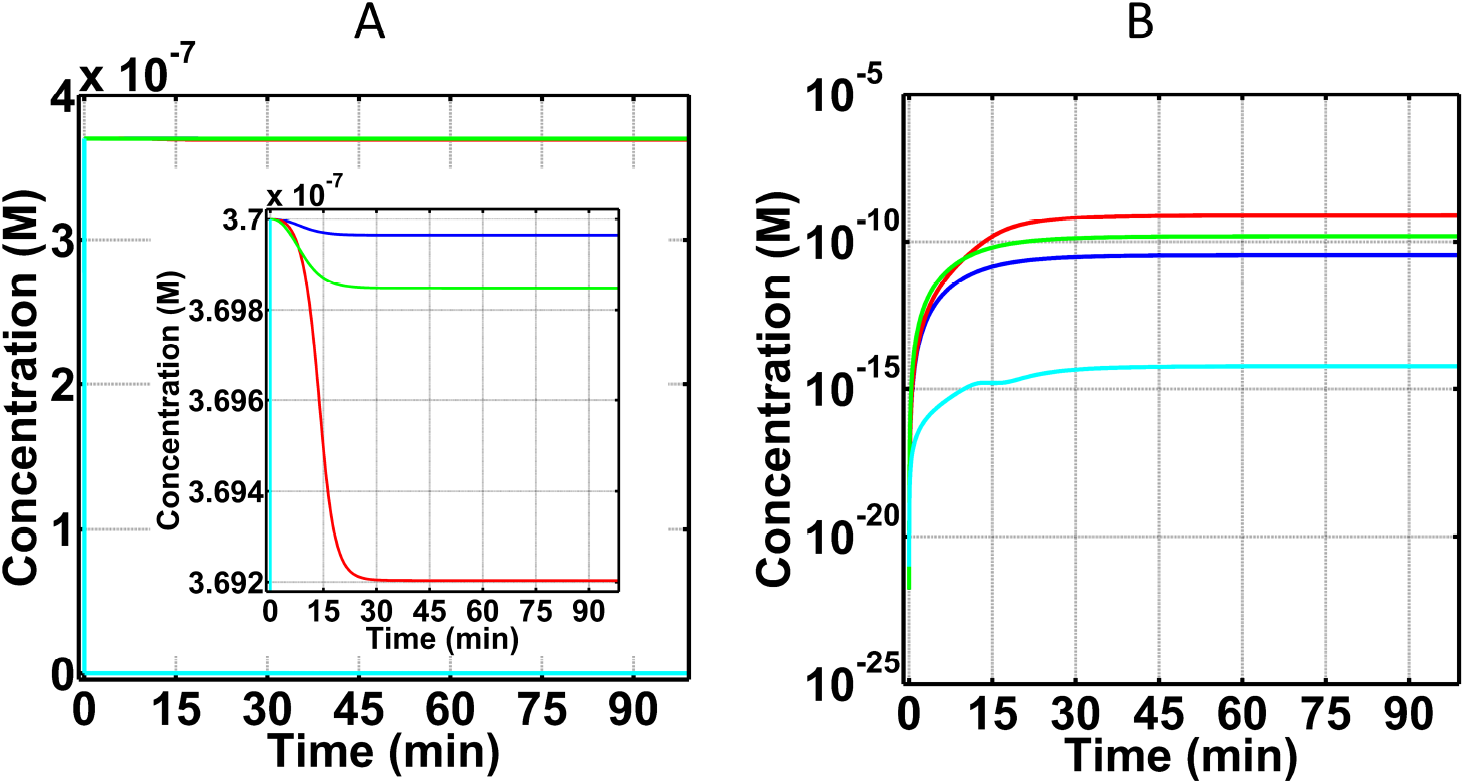
Concentration-time profiles for C5 and C5a-desArg under four conditions: (i) normal state (blue), (ii) FH disorder state (red), (iii) FH disorder state with compstatin treatment (green), and (iv) FH disorder state with eculizumab treatment (cyan). (A) Consumption of C5 under the four conditions. The inset is a zoom-in, showing that small amounts of C5 are consumed under the first three conditions (i–iii). Addition of eculizumab removed most of blood plasma C5 to form eculizumab:C5 complex (not shown). (B) Production of C5a-desArg after cleavage of C5. Treatment with eculizumab produced the lowest C5a-desArg levels, corresponding to about five orders of magnitude lower concentration compared with that of alternative pathway FH disorder.

#### 3.1.3. FB and its cleavage fragments, Ba and Bb, as biomarkers for FH disorder state

Complement FB plays a pivotal role in the consumption of C3 and C5 by providing the cleaving unit of the C3/C5 convertases. Once FB is in complex with C3b (C3bB), FD cleaves FB to form Ba and C3bBb. C3/C5 convertase C3bBb, however is very unstable and dissociates into C3b and Bb. In addition, complement regulators such as FH, CR1, and DAF can also increase the rate at which C3bBb dissociates. Figure 4 presents the time profiles of FB and its cleavage products Ba and Bb. In the normal state and FH disorder state with compstatin treatment, FB is reduced by < 2% relative to its blood plasma concentration (2.2× 10^−6^M). This minor reduction in FB highlights the regulatory role of FH when unimpaired but also the compensatory role of compstatin under compromised FH. However, in FH disorder state and FH disorder state with eculizumab treatment, the difference of FB relative to its blood plasma concentration is 48% and 49%, respectively. These values correspond to concentrations of 1.14×10^−6^M and 1.12×10^−6^M, respectively. Similarly, Figs 4B and 4C show time profiles for Ba and Bb where their levels are higher in FH disorder state and FH disorder state with eculizumab treatment. The concentrations of Ba and Bb reach values of ~1.05× 10^−6^M in FH disorder state, while Ba and Bb reach ~1.07×10^−6^M in FH disorder state with eculizumab treatment. In contrast, levels of Ba and Bb are about one order of magnitude lower under the normal state, reaching a concentration of 2.5 × 10^−8^M. Similarly, levels of Ba and Bb are reduced by about three orders of magnitude under FH impairment with compstatin treatment, reaching a concentration of 8.1 × 10^−10^M.

**Fig 4.**
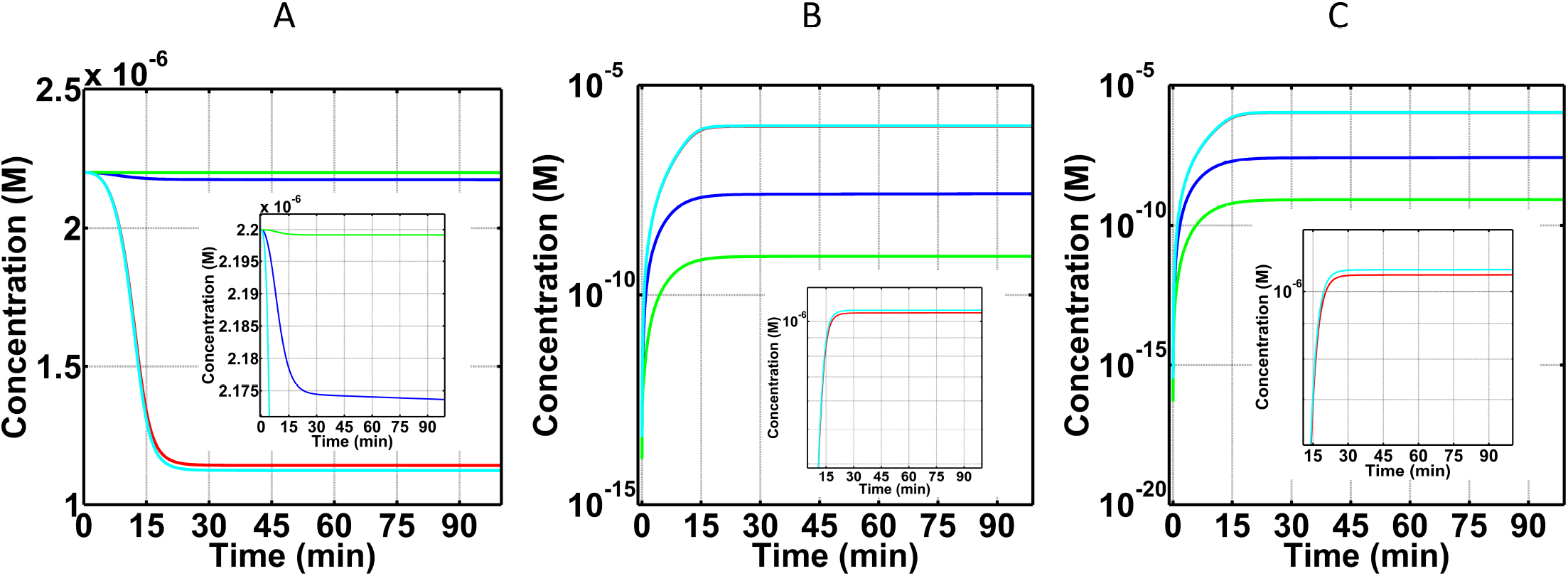
Concentration-time profiles for FB, Ba, and Bb under four conditions: (i) normal state (blue), (ii) FH disorder state (red), (iii) FH disorder state with compstatin treatment (green), and (iv) FH disorder state with eculizumab treatment (cyan). (A) FB shows the highest level of consumption in the FH disorder state and FH disorder state with eculizumab, demonstrating the minor effects of eculizumab on FB. On the contrary, compstatin nearly restores the concentration-time profile of FB to that of normal. (B and C) Similar, but opposite in magnitude effects for FB cleavage fragments Ba (Panel B) and Bb (Panel C). The FH disorder results to overproduction of Ba and Bb (view of the red graph is obscured by the cyan graph, see insets for distinction). Compstatin has a strong over-restorative effect (compare green with red and blue graphs), whereas eculizumab has a small effect (compare cyan with red and blue graphs). Insets are zoomed-in portions of the parent Panels, showing that FB levels are tightly controlled in normal state and FH disorder state with compstatin treatment (A), whereas Ba and Bb reach the highest production levels in the FH disorder state and FH disorder state with eculizumab treatment (B and C, respectively).

#### 3.1.4. Fluid phase MAC fC5b-9) as a biomarker for FH disorder state

After the cleavage of C5, a cascade of reactions ensues to form fluid phase C5b-9 (fC5b-9 or fMAC). As shown in Fig 5, the time profiles of fC5b-9 follow similar trends as C5a-desArg. The highest concentration of fC5b-9 (8.2× 10^−16^M) was in FH disorder state, while the FH disorder state with compstatin treatment generated a lower fC5b-9 concentration of 1.5×10^−16^M. The normal state produced about one order of magnitude lower of fC5b-9, with concentration 3.7× 10^−17^M, relative to FH disorder state. However, in FH disorder state with eculizumab treatment, fC5b-9 levels are about four orders of magnitude lower (7.7×10^−21^M) relative to those in FH disorder state with compstatin treatment.

**Fig 5.**
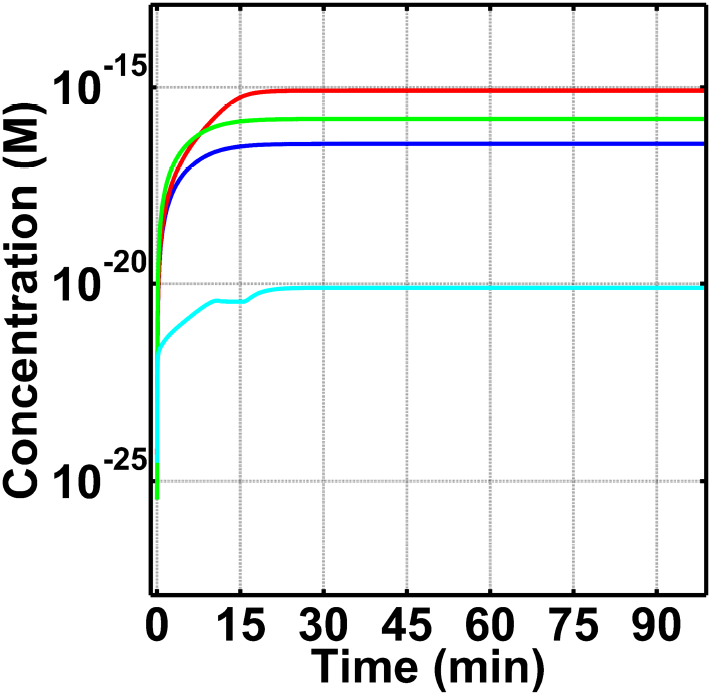
Concentration-time profiles for fC5b-9 under four conditions: (i) normal state (blue), (ii) FH disorder state (red), (iii) FH disorder state with compstatin treatment (green), and (iv) FH disorder state with eculizumab treatment (cyan). Highest concentration of fC5b-9 is produced in FH disorder state, with compstatin showing a restorative trend, but incomplete restoration, and eculizumab showing an over-restorative effect. Production of fC5b-9 under FH disorder is about one order of magnitude higher than normal. Compstatin treatment generates lower levels of fC5b-9 than FH disorder state, but fC5b-9 levels are still higher compared to normal state. Finally, eculizumab treatment shows production of fC5b-9 to be about three orders of magnitude lower than normal.

### 3.2. Sensitivity analysis

Global sensitivity analysis is performed using multi-parametric sensitivity analysis (MPSA) to identify critical parameters that affect the central complement substrate C3 and the terminal substrate C5 under normal condition. The results are shown in Figs 6A and 6B. C3 levels are predominately affected by classical pathway activation and propagation in the fluid state. The most sensitive parameter is the one that governs the interaction between regulator C1-INH and activated fluid phase C1*, as shown in Fig 6A. This is followed by other classical pathway regulation steps between C4BP:fC4b, decay of C3 convertase (C4bC2a:C4BP), and enzymatic parameters (k_cat_ and K_m_) of C1* targeting C4bC2. Similarly, the enhancement of the terminal cascade through the classical pathway is seen for the key interactions that affect levels of C5 (Fig 6B). This includes the activation step of C5 through the k_cat_ of C3bC4bC2a, dissociation rate constant of C3bC4bC2a, and the regulatory steps of C4BP with fC4b and C3/C5 convertase (C4bC2a). Lastly, key interactions of the alternative pathway that affect the levels of C3 and C5 involve the inactivation step of C3b in complex with FH (C3bH). This complex is targeted by FI to form iC3b. Figures 6A and 6B show that inactivation of C3b in C3bH by the action of FI is an important step in mediating the levels of C3 and C5.

**Fig 6.**
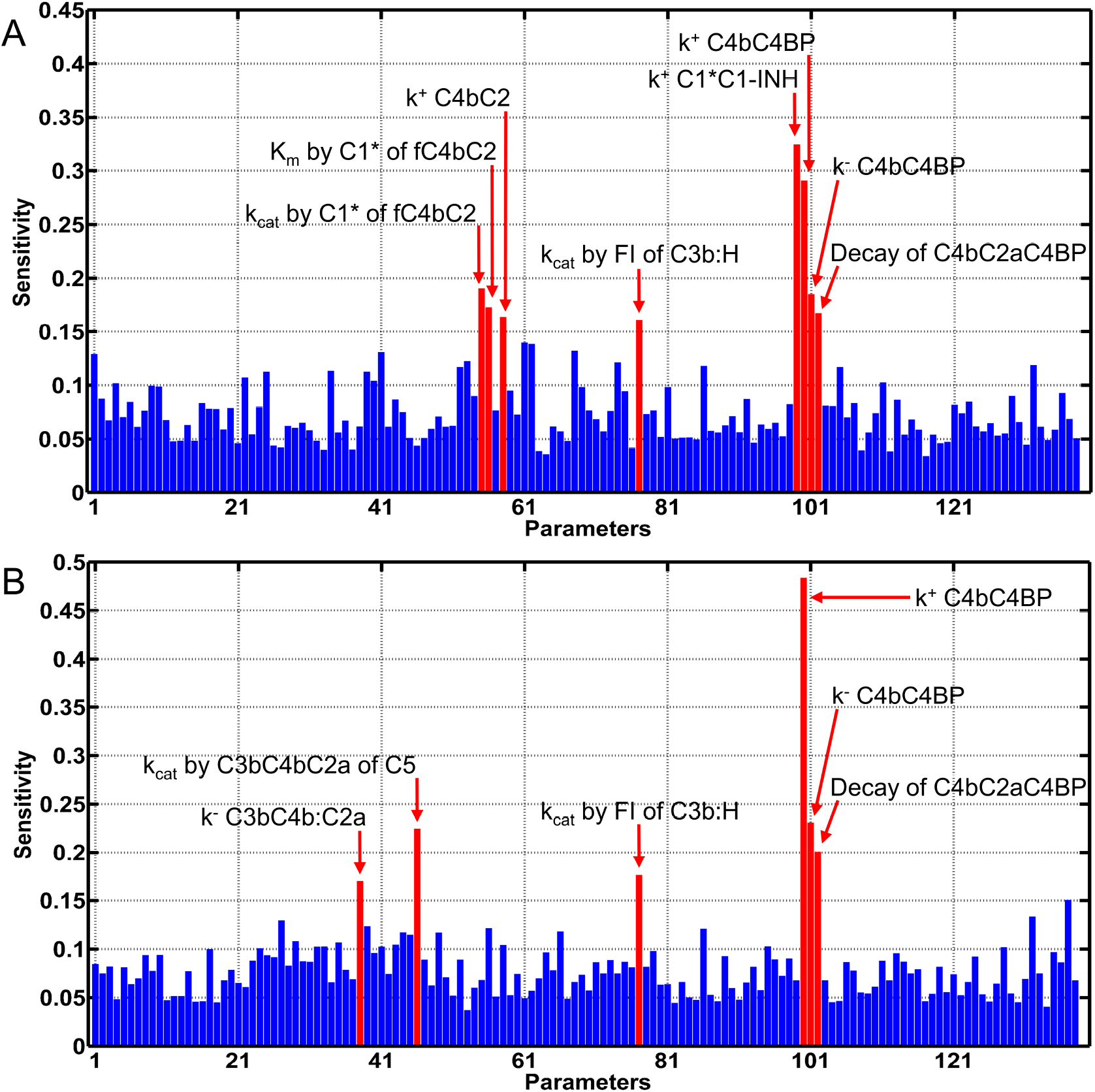
Global sensitivity analysis. Multi-parametric sensitivity analysis was performed to identify parameters that affect central substrates C3 and C5 under normal conditions. Both substrates are sensitive to the activation and propagation of the classical pathway. (A) Regulation of C1* through the action of C1-INH is the most sensitivity parameter that mediates C3 levels. (B) C5 levels are predominantly affected by regulation of C4b through C4BP. This step inhibits formation of C3/C5 convertases. In addition, both C3 and C5 are affected by the alternative pathway regulation step that inactivates C3b through the actions of FH in conjunction with FI (Panels A and B).

### 3.3. Dose dependence analysis for compstatin and eculizumab in modulating FH disorder state

Our results thus far have shown that using compstatin and eculizumab with a concentration of 20-fold higher than the concentration of their respective targets, C3 and C5, to be more than adequate in modulating complement biomarkers in the FH disorder state. Furthermore, these concentrations outperformed complement regulators under normal conditions by generating lower levels of complement biomarkers such as C3a-desArg, Ba, Bb, and fC5b-9 (Figs 2B, 4, and 5). Because of this, we consider the concentration of compstatin of 1.4 × 10^−4^M (20-fold higher than C3), and the concentration of eculizumab of 7.4 × 10^−6^M (20-fold higher than C5), to be upper bounds, and investigated the effect of drug lower doses on complement biomarkers. We use C3a-desArg as early-stage biomarker and fC5b-9 as late-stage biomarker to assess the effects of the two drugs in complement modulation. Figures S1-S6 of the Supplementary Material show the concentration dependences of the remaining biomarkers, C3, C5, C5a-desArg, FB, Ba, and Bb.

Figure 7 shows concentration-time profiles for C3a-desArg in the FH disorder state with compstatin concentrations of 7.1×10^−6^M (one-to-one to C3; Fig 7A), and 1.4×10^−6^M (5-fold lower than C3; Fig 7B), for comparison to Fig 2B (compstatin concentrations of 1.4× 10^−4^M, 20fold higher than C3). The same fold-change in concentration was used for eculizumab: 3.7× 10^−7^M (one-to-one to C5; Fig 7A), and 7.4× 10^−8^M (5-fold lower than C5; Fig 7B), for comparison to Fig 2B (eculizumab concentration of 7.4×10^−6^M, 20-fold higher than C5). The results show that the effect of one-to-one compstatin-C3 concentration nearly recovers the C3a-desArg concentration-time profile to that of normal state (Fig 7A), whereas 5-fold lower compstatin concentration than C3 concentration is insufficient to regulate (Fig 7B). On the other hand, the effect of 20-fold higher compstatin concentration than C3 concentration is over-regulating (or over-restorative), keeping the C3a-desArg levels lower than those of the normal state (Fig 2B). Lastly, variations in eculizumab have minor effects on regulating early complement fragment C3a-desArg (Figs 7A and B).

**Fig 7.**
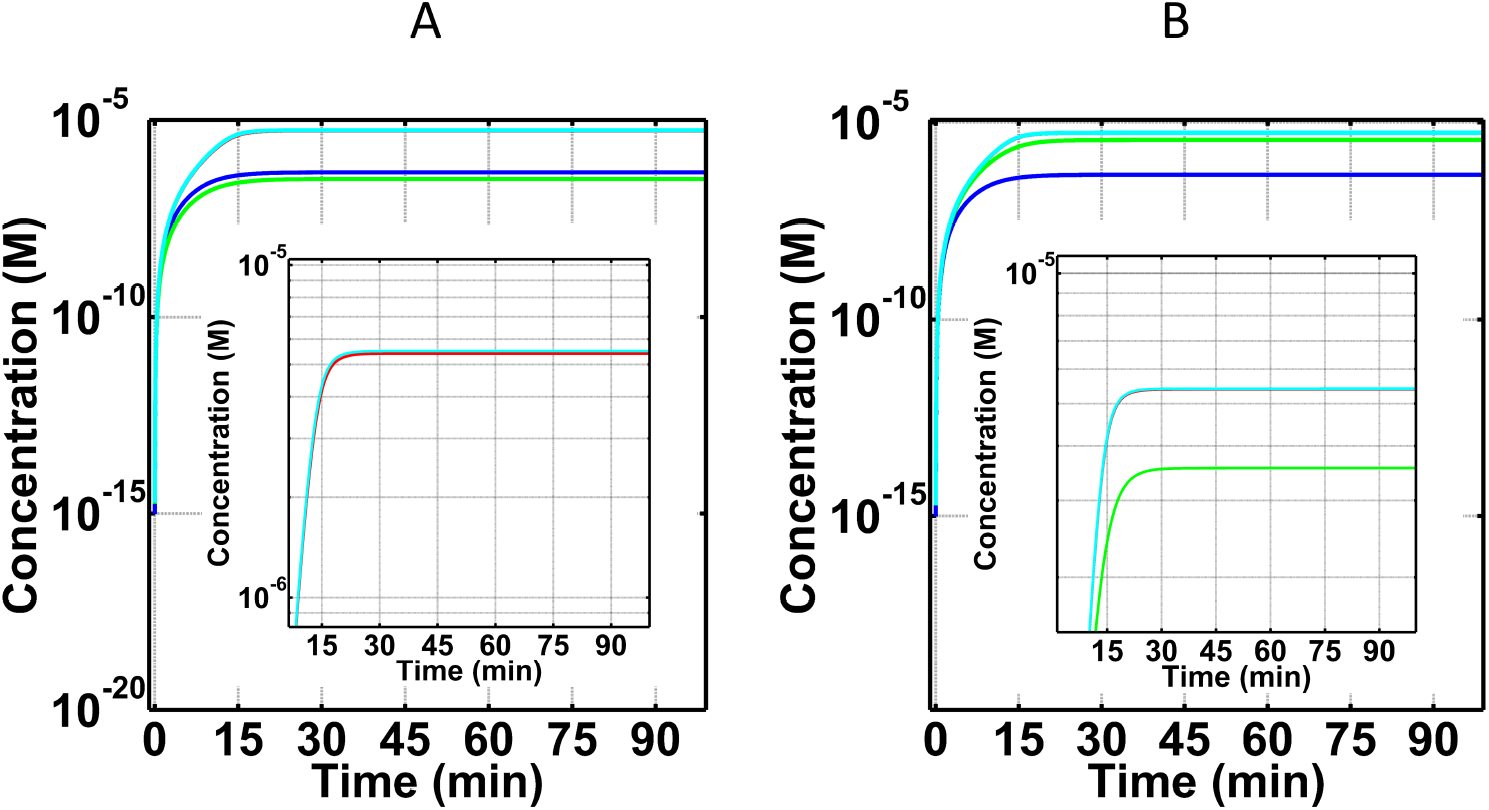
Concentration-time profiles for C3a-desArg under four conditions: (i) normal state (blue), (ii) FH disorder state (red), (iii) FH disorder state with compstatin treatment (green), and (iv) FH disorder state with eculizumab treatment (cyan). Concentrations of compstatin used: 7.1 × 10^−6^M (one-to-one with C3; A), and 1.4× 10^−6^M (5-fold lower than that of C3; B). Concentrations of eculizumab used: 3.7×10^−7^M (one-to-one with C5; A), and 7.4×10^−8^M (5-fold lower concentration than that of C5; B). A dose-dependent response is observed for C3a-desArg under different concentrations of compstatin. Low levels of C3a-desArg are produced under the higher compstatin concentration of 7.1 × 10^−6^M (A), whereas high levels of C3a-desArg are produced under the lowest compstatin concentration of 1.4× 10^−6^M (B). Restorative effects of compstatin are compromised at a dose of 5-fold lower concentration than the concentration of target protein C3. Conversely, varying concentrations of eculizumab had minor effects on the levels of C3a-desArg, as shown in the insets of Panels (A and B).

After examining the effects of varying the concentrations of compstatin and eculizumab on C3a-desArg, we next investigated the effects of these inhibitors on late stage biomarker fC5b-9 (Fig 8). We used the same variations in concentrations for compstatin and eculizumab as used for C3a-desArg, described above. Different doses of compstatin had small effect and never recovered the fC5b-9 concentration-time profile to that of the normal state (Figs 5 and 8). In contrast, one-to-one eculizumab-C5 concentration nearly recovers the fC5b-9 concentration-time profile to that of normal state (Fig 8A), whereas 5-fold lower eculizumab concentration than C5 concentration is insufficient to regulate (Fig 8B). As discussed above, the effect of 20-fold higher eculizumab concentration than C5 concentration is over-regulating (or over-restorative), keeping the fC5b-9 levels lower than those of the normal state (Fig 5).

**Fig 8.**
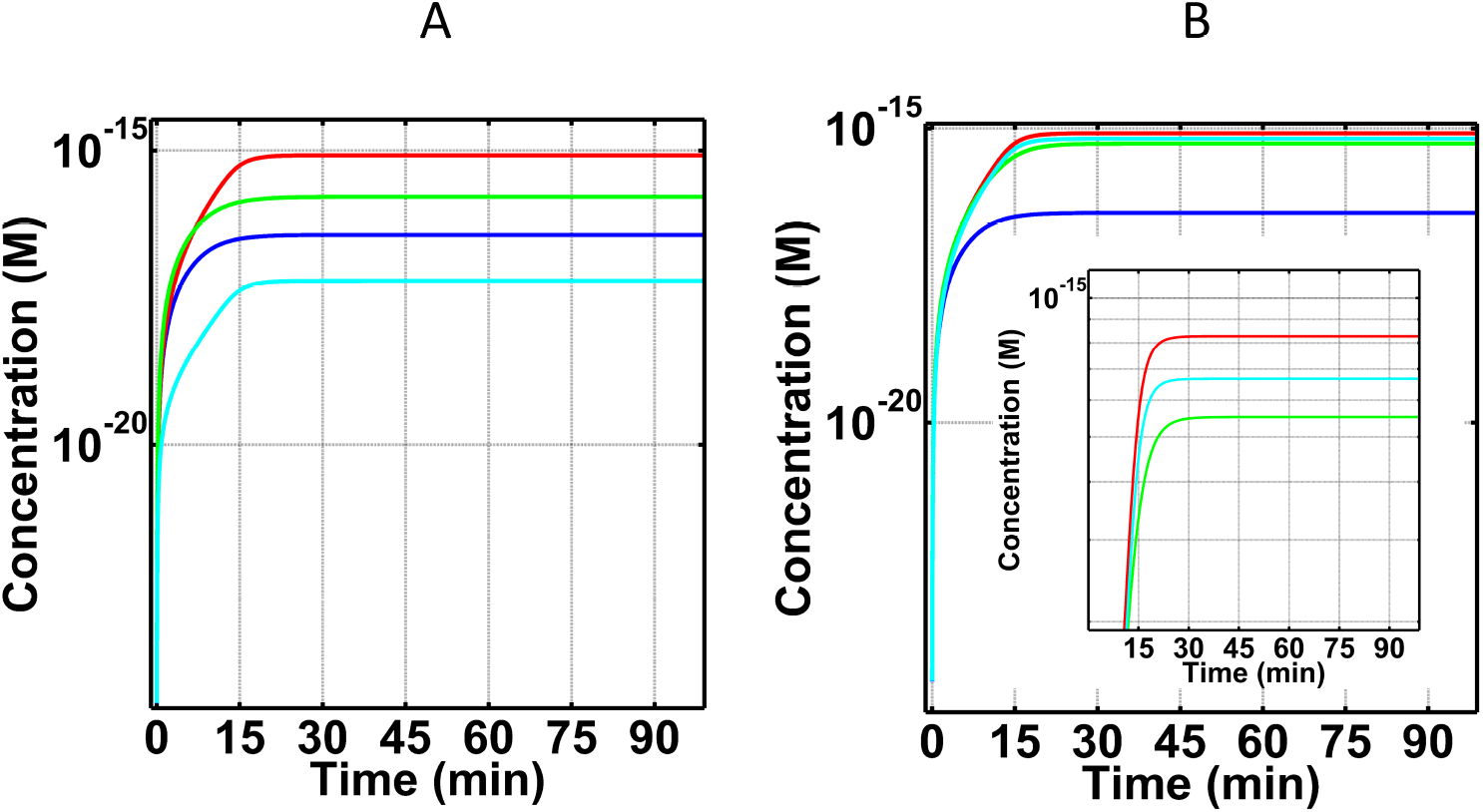
Concentration-time profiles for fC5b-9 under four conditions (i) normal state (blue), (ii) FH disorder state (red), (iii) FH disorder state with compstatin treatment (green), and (iv) FH disorder state with eculizumab treatment (cyan). Concentrations of compstatin used: 7.1 × 10^−6^M (one-to-one with C3; A), and 1.4× 10^−6^M (5-fold lower than that of C3; B). Concentrations for eculizumab used: 3.7×10^−7^M (one-to-one with C5; A), and 7.4×10^−8^M (5-fold lower than C5; B). Treatment with different doses of compstatin showed small effects on fC5b-9 levels. In contrast, treatment with varying concentrations of eculizumab generated a dose-dependent response for fC5b-9. Lowest fC5b-9 levels are produced under the highest eculizumab dosage, and vice versa. Similar to the insufficient dose of compstatin in restoring C3a-desArg, eculizumab also loses restorative effects if the used dose is 5-fold lower than the concentration of the target protein C5.

## 4. Discussion

### 4.1 Modeling of fluid phase alternative and classical pathway activation, and relation to FH-based diseases

In the present study, we used mathematical modeling of the complement system to understand the complex interplay between complement activation/propagation and regulation. First, differential equations were used to model the biochemical reactions involved in the alternative and classical pathways of the complement system under normal condition (physiological condition in homeostasis). Second, we modeled an alternative pathway disordered state by impairing FH to better understand the dynamics of complement pathology in FH-mediated diseases, called herein FH disorder state. This negative regulator, FH, was chosen because alternative pathway-associated disorders, such as AMD, aHUS, C3GN, DDD, implicate FH impairment at the levels of lower effective concentration or altered binding kinetic parameters leading to improper regulation. Third, we added two complement therapeutic states by incorporating FH disorder state with compstatin treatment (targeting C3) and FH disorder state with eculizumab treatment (targeting C5). And lastly, using these four computational states, we generated concentration-time profiles for biomarkers associated with alternative pathway dysregulation, such as C3, C3a-desArg, C5, C5a-desArg, FB, Ba, Bb, and fC5b-9.

### 4.2. Biomarkers under normal state

Our results show only 7% consumption of C3 (Fig 2A) and <1% consumption of C5 (Fig 3A) relative to their respective starting blood plasma concentrations, under normal state. This low consumption reflects the potent regulatory effect of a physiological (unperturbed) FH has on the system. For instance, FH can induce regulation on early phases of the alternative pathway by targeting and inactivating (in conjunction with FI) complement components such as C3(H_2_O) and C3b. In addition, FH can regulate enzymes C3(H_2_O)Bb and C3bBb, by accelerating their decay rates and inhibiting propagation of complement in the fluid and surface phases. Moreover, another indicator of proper regulations is shown on the levels of FB consumption and generation of its cleavage product Ba and Bb. Binding of FB to either C3(H_2_O) or C3b will form the proconvertases of alternative pathway that later become activated through FD to form C3(H_2_O)Bb and C3bBb. However, presence of FH modulates early complement species to ensure most of plasma FB will not be involved in either forming proconvertases or activated through cleavage. The results of Fig 4A reflect these regulatory events where small amounts of FB are consumed in the process of forming short-lived alternative pathway convertases and cleavage fragments of FB, Ba (after cleavage by FD; Fig 4B) and Bb (after convertase dissociation by FH, FHL, and CR1, or self-dissociation; Fig 4C), are also produced in small amounts. Subsequently, proper regulation of convertases also leads to proper modulation of the cleavage rates of C3 and C5. This results in lower amounts of C3 being converted to nC3b and C3a (further deactivated to C3a-desArg), and C5 into C5b and C5a (further deactivated to C5a-desArg). Our model shows these events where the production of biomarkers C3a-desArg (Fig 2B) and C5a-desArg (Fig 3B) are produced in low amounts under normal conditions. However, C3a-desArg is shown to be at a higher concentration than C5a-desArg. And lastly, a termination step of the complement system is instigated by C5b associating with C6, C7, C8, and C9 to form fluid phase MAC/fC5b-9. Since the conversion of C5 into C5b and C5a is properly modulated, small amounts of fC5b-9 are produced as a result in the normal state (Fig 5).

### 4.3. Comparison of normal and FH disorder states

Introduction of alternative pathway dysregulation, through FH impairment, referred herein as FH disorder state, significantly reduced C3 levels by 76% relative to C3 blood plasma concentration (Fig 2A). This means that suboptimal FH function results in increased levels and stability of alternative pathway C3/C5 convertases. These enzymes rapidly consume blood plasma C3 to produce elevated levels of C3a-desArg (after C3a deactivation; Fig 2B) and nC3b compared with the normal state. Presence of elevated nC3b subsequently forms the basis for more production of convertases that later enhance the propagation step of alternative pathway by not only cleaving more C3, but also C5. Figures 3A and 3B exemplify this event where more C5 is consumed, and C5a-desArg (after C5a deactivation) levels are elevated under the alternative pathway dysregulation of the FH disorder state than in the normal state. Additionally, levels of plasma C5b rise as more C5 is consumed. This initiates the terminal step by C5b starting a cascade of reactions to form fluid phase MAC/fC5b-9. In comparison to the normal state, elevated levels of fC5b-9 are generated under FH disordered state (Fig 5).

Altogether these data show that impairing FH has the effect of compromising the regulatory checkpoints to generate an overly active complement system with reduced levels of C3 and C5, coupled with elevated levels of inflammation markers C3a-desArg and C5a-desArg.

### 4.4. Effect of compstatin on FH disorder state

Compstatin inhibits cleavage of C3 to C3a and nC3b, and thus inhibits C3 consumption. The rapid reduction of C3 concentration shown in Fig 2A is not due to consumption, but rather it is owed to C3 being in complex with compstatin to form compstatin:C3 complexes (not shown in Fig 2). Compstatin has an over-restorative effect on C3a-desArg, meaning that it brings down the concentration of C3a-desArg in the FH disorder state by about three orders of magnitude, while C3a-desArg levels in the normal state are about one order of magnitude lower than the FH disorder state. In contrast, compstatin has an under-restorative effect on C5a-desArg, meaning that although it brings down the concentration of C5a-desArg in the FH disorder state, it is still about one order of magnitude higher than the C5a-desArg concentration under the normal state (Fig 3B).

In summary, an overly active complement system under compstatin treatment better modulates levels of C3a-desArg than C5a-desArg. This outcome is due to regulation induced by compstatin in the early phases of complement activation to generate lower amounts of C3a, later becoming deactivated to form C3a-desArg. Although there is a compensatory effect, it is not clear what is the net effect of compstatin inhibition on C3a-/C5a-induced inflammatory response, given that C5a is a more potent pro-inflammatory protein than C3a. It has been suggested that C3a has both proinflammatory and anti-inflammatory properties [50], whereas C5a is proinflammatory. Therefore, in certain pathologies, C3a and C5a may have opposite roles, and this should be taken into account when selecting treatment drugs.

Furthermore, treatment with compstatin generates a trend for fC5b-9 that is similar to C5a-desArg but smaller in concentration magnitude (Fig 5). This is because the formation of fC5b-9 is based on the other C5 cleavage product, C5b, which is equivalent in concentration to C5a immediately after cleavage; however C5b is further involved in biochemical reactions that include inactivation, propagation on host cells, and regulation in fluid and host cells (Fig 1).

In terms of FB and its cleavage fragments (Ba and Bb), compstatin has nearly complete restorative effect on FB, and over-restorative effects on Ba and Bb. These results highlight on the compensatory role that compstatin plays in cases of FH impairment. The potency of FH lies on its ability to target early fragments of the complement activation (C3b) and enzymes (C3/C5 convertases) which initiate and extend the propagation step of the alternative pathway. But in cases of FH impairment, compstatin can aid complement regulation by first targeting C3. This results in reduced levels of early complement fragment C3b. In addition, since initiation and extension of the propagation step depends on C3b and FB, the presence of compstatin reduces C3b levels and this reduces the rate of FB consumption and generation of its cleavage products (Ba and Bb) as shown in Fig 4.

### 4.5. Effect of eculizumab on FH disorder state

Eculizumab has a minor effect on C3 and its cleavage fragments, C3a and nC3b (Fig 2), and on FB and its cleavage fragments, Ba and Bb (Fig 4). This is because eculizumab acts on C5, and inhibits cleavage of C5 to C5a and C5b, thus inhibiting C5 consumption. The dramatic reduction of C5 concentration shown in Fig 3A is not due to consumption, but rather because treatment with eculizumab generates eculizumab:C5 complexes (not shown in Fig 3). Eculizumab over-restores the concentrations of C5a-desArg and fC5b-9 in the FH disorder state to about four and three, respectively, orders of magnitude lower than the C5a-desArg and fC5b-9 concentrations in the normal state.

These results suggest that eculizumab may be a better inhibitor than compstatin for diseases that are driven by excess MAC/C5b-9 formation. The potency of eculizumab as a terminal cascade inhibitor is shown on the levels C5a-desArg and fC5b-9 concentrations. However, even in the absence of compstatin and eculizumab, in the FH disorder state the change in C5 concentration is small compared with the change in C3 concentration, relative to their concentration changes in the normal state. Depending on the disease, a treatment with inhibition at the early stages of the complement pathway (e.g. C3), or a treatment at the late stages of complement pathway (e.g. C5), or a dual treatment may be necessary. The model may be useful to select the point of inhibition, at C3, C5, or other (e.g. FB, FD, fC5b-9, etc), depending on the specific effect we aim to suppress, e.g. inflammation, opsonophagocytosis, MAC/C5b-9 formation.

### 4.6. Sensitivity analysis

Our sensitivity analysis showed the classical pathway fluid phase activation as a major cross activator of the alternative pathway (Fig 6) under normal conditions. The alternative pathway is initiated through tick-over process with a first order kinetic rate constant of 4.5× 10^−6^ s^-1^ while the classical pathway is spontaneously activated much faster through C1 with a kinetic rate constant of 2.8×10^−3^ s^-1^ [4,51]. Activated C1 (C1*) cleaves C4, followed by C4bC2 to form the classical pathway C3/C5 convertase C4bC2a. This enzyme can initiate the alternative pathway by cleaving C3 before the tick-over forms the initial convertase of the alternative pathway, C3(H_2_O)Bb. Thus, having a faster activation rate makes C1 the critical step in forming enzymes that can activate and propagate the alternative pathway in the fluid phase. Figure 6A displays this notion where the kinetic parameter that regulates C1* by the action of C1-INH is the most sensitive parameter that mediates levels of C3. Lastly, while the remaining critical parameters of the classical pathway are involved in interactions that affect steps that either form or inhibit C3/C5 convertases of the classical pathway, the alternative pathway has a single key regulation step mediated by FH in conjunction with FI that controls the levels of C3 and C5 (Fig 6). The potency of FH not only lies in accelerating the decay rate of convertases, but also working in concert with FI to cleave C3b into iC3b. This cleavage step reduces the rate of C3/C5 convertases formation since iC3b does not from convertases.

### 4.7. Effects of varying doses of compstatin and eculizumab on C3a-desArg and fC5b-9 in FH disorder state

Varying concentrations of compstatin affect levels of C3a-desArg in a dose-dependent manner, while different doses of eculizumab showed minor effects on C3a-desArg (Figs 2B and 7). Unlike eculizumab, treatment with compstatin generate compstatin:C3 complexes that are shielded from cleavage by C3/C5 convertases. This step inhibits early cascade of reactions induced by nC3b and C3a that later form C3/C5 convertases and C3a-desArg, respectively. However, reducing the levels of compstatin further compromises regulation and increase the levels of C3a-desArg. For instance, restorative capabilities of compstatin are significantly impaired when the concentration of compstatin is 5-fold lower than that of C3. Under this concentration, compstatin has an under-restorative effect by generating C3a-desArg levels that are closer to the FH disorder state than the normal state.

Although production of the early-stage complement biomarker C3a-desArg is highly dependent on compstatin and not eculizumab, late-stage complement fC5b-9 is more dependent on eculizumab than compstatin (Figs 5 and 8). Varying the doses of compstatin generated fC5b-9 levels that were closer to fC5b-9 levels under the FH disorder state. However, treatment with eculizumab generates a dose-dependent response of fC5b-9 levels because of the eculizumab:C5 complexes that are formed. This eculizumab:C5 complex is protected from cleavage by convertases to C5a/C5b fragments, and thus C5b-based initiation of the cascade of reactions that lead to fC5b-9 formation is inhibited. As the levels of eculizumab increase, less fC5b-9 complexes are formed. Comparison of the results of Figs 5 and 8 show this relation where fC5b-9 levels are the lowest under the highest eculizumab concentration (20-fold higher than the concentration of C5). Conversely, highest levels of fC5b-9 are generated under the lowest eculizumab concentration (5-fold lower than the concentration of C5).

In summary, compstatin shows higher restorative efficacy for early-stage biomarker C3a-desArg, whereas eculizumab has higher restorative efficacy for late-stage biomarker fC5b-9. Also, one-to-one drug-to-target protein concentrations are sufficient for complete restoration of the FH disorder state to the normal state in this model. However, we should point out that this concentration ratio may change when we incorporate in the model drug metabolism and complement protein turnover steps.

### 4.7. Relation of the model to clinical disorders and therapeutics

Altogether our data highlight the importance of FH in regulating the levels of the central substrate of the complement system, C3. Furthermore, the presence of unimpaired FH is also essential for regulating the levels of blood plasma FB. Disorders related to the alternative pathway, such as C3GN, DDD, and aHUS are typically characterized with low levels of plasma C3 and FB, while FB cleavage products, Ba and Bb, are elevated [23–27]. Although these disorders are also associated with mutations/polymorphisms in other complement protein, such as C3, FI, FB, and FH related proteins, in addition to FH mutations/polymorphisms, our data shows that just impairing FH is sufficient to generate clinically observed trends for C3, FB, Ba, and Bb. In support of this notion, biomarker profiling of patients with C3GN and DDD shows that having type 1 mutation in FH (reduced levels of FH concentration) was sufficient to generate to low levels of plasma C3 and FB [27]. Furthermore, absence of plasma FH has also been observed to cause C3 glomerulopathies in humans [27]. Thus, FH plays a pivotal role in ensuring proper regulation, and mutations that alter concentration or kinetics of FH will instigate the dysregulation of the alternative pathway.

In addition to generating biomarker trends associated with the alternative pathway dysregulation, our computational model also shows treatment with eculizumab effectively regulates terminal complement activity while having a minor effect on early complement activity induced by C3. This coupled effect was observed for paroxysmal nocturnal hemoglobinuria (PNH) patients where treatment with eculizumab regulated terminal complement activation but some patients still suffered from extravascular opsonophagocytosis (also referred to as extravascular hemolysis) that is attributed to early phase complement activation and propagation induced by C3 [52]. Subsequently, a recent study showed that complement intervention by compstatin protected PNH erythrocytes from complement mediated lysis and modulated early phase complement deposition and propagation induced by C3 [53]. These findings are consistent with our computational model where eculizumab shows little effect on C3 level under FH impairment, whereas compstatin regulates potently early-stage complement activity and less potently late-stage complement activation. However, our model also shows eculizumab is much more potent regulator of late-stage complement activation and propagation than compstatin. Thus, efficient complement therapy may require disease-specific biomarker targeting, or dual point targeting.

Overall, this study serves as a proof-of-concept on the capabilities of the model in predicting the dynamics of the complement system under homeostasis and disease states, and in predicting the effects of drugs in restoring the complement system from a disease state to the normal state.

## Conclusion

We have generated a comprehensive model of the alternative and classical pathway of the complement system that is divided into: (i) normal state, corresponding to homeostasis in a healthy person, (ii) alternative pathway dysregulation through FH impairment, called FH disorder state, (iii) FH disorder state with compstatin treatment, and (iv) FH disorder state with eculizumab treatment. We analyzed the state of the system by generating time profiles for biomarkers associated with alternative pathway FH disorders: C3, C3a-desArg, C5, C5a-desArg, FB, Ba, Bb, and fC5b-9. The computational model shows major changes on these biomarkers in the FH disorder state, compared with the normal state. The model shows restorative effects in the biomarker concentration levels, from the FH disorder state towards the normal state, upon treatment with compstatin and eculizumab. Compstatin is more effective in restoring biomarker concentrations at the early stages of the complement cascade, compared with biomarkers of the late stages. Conversely, eculizumab is more effective in restoring concentrations at the late stages of the complement cascade, compared to the effects of compstatin on late stage biomarkers. Lastly, the model predicts effective doses of compstatin and eculizumab to be one-to-one against their respective targets, C3 and C5, to properly restore early- and late-stage biomarkers.

The model serves as the basis for developing disease-specific models, and even patient-specific models if sufficient genetic and clinical data is available, by perturbing other complement proteins (than or in addition to FH), or combinations of proteins. The model can be used to identify the optimal point of inhibition within the complement cascade, and if a pool of drugs is available, the model can be used to select the right drug or combination of drugs for treatment. Currently, eculizumab is approved for use in the clinic, and compstatin is undergoing clinical trials. There other approved drugs targeting C1-INH, and there are several other drugs in the pre-clinical and clinical trial pipeline, targeting C1r, C1s, C3, C3a, C5, C5a, C6, FH, FD, FB, FP, etc [54] and soon we expect to have approval for clinical use a pool of several drugs against complement-mediated inflammatory and autoimmune diseases.

## Acknowledgements

We thank Rohaine Hsu for helping with the detailed inspection of the equations of the computational model.

## Appendix A. Supplementary data

Supplementary data associated with this article can be found, in the online version, at http://….

